# CVAE-based Causal Representation Learning from Retinal Fundus Images for Age Related Macular Degeneration(AMD) Prediction

**DOI:** 10.1101/2025.02.13.638092

**Authors:** Daeyoung Kim

**Affiliations:** The Department of Applied Statistics, Yonsei University, Seoul, South Korea

**Keywords:** Age-related macular degeneration, Causal representation learning, Computer vision, Retinal disease, Neural Causal Model, VAE

## Abstract

Regarding relatively poor prognosis and acute vision impairment, analyzing Age-Related Macular Degeneration, or AMD has been one of the most important tasks in retinal disease analysis. Especially, constructing methods to analyze and predict Wet AMD, which is characterized by rapid RPE damage due to neovascularization, has been a demanding task for many ophthalmologists for decades. Recently, with advancements in ML/DL frameworks and computer vision AI, these previous efforts are now leading to drastic enhancements in AMD prediction and mechanism analysis. Specifically, use of attention mechanism based CNNs or XAI methods are leading to higher performance in predicting AMD status and reliable explanations. Under current success in the usage of cutting-edge techniques, this research implemented a novel latent causal representation learning framework to further enhance AI-based models to comprehend complex causal AMD mechanisms with only access to retinal fundus images, while constructing a more reliable type of AMD prediction model. Results show that valid convolutional VAE and GAE based explicit latent causal modeling can lead to successful causal disentanglements of underlying AMD mechanisms, while returning essential causal factors that can be utilized to reliably distinguish normal fundus and AMD fundus images in downstream tasks such as diagnosis prediction.

## I. Introduction

**H**aving relatively poor prognosis, Age-Related Macular Degeneration(AMD or ARMD) has been a significant threat to the human vision along with a variety of retinal diseases such as Glaucoma or Diabetic Retinopathy [1,2]. Especially among retinal damage, Wet AMD has been one of the most major causes of acute vision impairment, which rapidly leads to complete or permanent loss of vision. Aside from having severe prognosis, AMD is also getting substantial attention for its increase in scope. According to the CDC, while being the primary cause of permanent vision impairment in people who are aged 65 and older, it was found that approximately 18.34 million in the US who are aged 40 and older suffered from early-stage AMD in 2019 [3,4]. In global aspects, increase rate of AMD patients is expected to be 47% in 2040 (from 196 million in 2020 to 288 million in 2040) [5]. Due to its current severity and linkage to mental health [6], numerous researchers probed for methods to enhance the possibility of earlier AMD detection or analyzed mechanisms of AMD-based vision loss to find effective treatments. In traditional works, manual OCT/FA image interpretations to track optical components (such as RPE or photoreceptors) and biomarker-based risk evaluations were mainly implemented for AMD analysis [7-10]. However, due to the intrinsic complexity in AMD pathogenesis and a broad overlap between other dystrophic macular disorder features, accurate or timely diagnosis of AMD still remained as a demanding task [11].

In recent works, with advancements in machine learning and neural network analysis, the possibility of overcoming these limitations is being increased in various aspects. Under extreme advancements in artificial intelligence, ML/DL based modelling approaches show high performance in fulfilling previous objectives. Specifically, methodologies such as attention mechanism based Convolutional Neural Networks (CNN) for retinal image classification or XAI-based analysis for acute image classifications are repeatedly being implemented for DL-based AMD analysis, leading to substantial advancements in retinal disease recognition or disease prediction performance [12-14]. Furthermore, fast developments in deep generative AI have enabled researchers to generate synthetic retinal images, which led to precise recognition or comprehension regarding optical components.

Based on the success in numerous applications of DL/ML methods in retinal analysis, this study combined cutting edge techniques in deep learning with AMD analysis to probe for novel mechanisms that can lead to accurate AMD diagnosis and valid interventional analysis in the aspect of treatments under better reliability. To be specific, this study attempted to extract latent causal representations related to AMD diagnosis from patient’s retinal fundus images using Convolutional VAE and Neural Causal DAG extraction. Unlike previous works, which only focused on correlation or classification performance, this research also focused on making computer vision models comprehend the causal mechanism of AMD and return valid inference regarding interventions (treatments), while achieving high prediction performance regarding AMD development.

## II. Preliminaries

### A. AMD mechanism as domain knowledge

To check success in causal disentanglements under latent causal representations extracted from retinal fundus images, a basic empirical causal graph, which can be deduced from domain knowledge of AMD development, was set as a benchmark. AMD diagnosis, or vision impairment due to AMD is a result of either two distinct processes: Dry AMD and Wet AMD [15,16]. Characterized by atrophy in RPE, Dry AMD is resulted from cumulation of drusen, an undisposed metabolic waste. With increased oxidative stress, proteins and lipids are abnormally deposited under the retina as specks. These specks, or drusen, disturb photoreceptors from receiving nourishment as it hinders interaction between blood vessels and the retina [16,17]. This not only leads to deterioration of RPE cells but also vision impairment as the size of drusen gets larger.

On the other hand, in Wet AMD, which is characterized by hemorrhage or intraretinal fluids within the retina, neovascularization is considered as a major cause. Due to structural changes in choroidal vasculature from abnormal vascular endothelial growth factor(VEGF) levels, angiogenesis or neovascularization occurs in Wet AMD, which leads to damage in RPE, bruch’s membrane and photoreceptors. This leads to generation of intraretinal fluids and hemorrhage which may result in focal loss and vision impairment by damaging the macula itself [15]. Thus, AMD mechanisms can be summarized into a simple causal diagram as in Fig 1. (Though there were some experimental results where possibility of interactions between drusen and neovascularization was found [17,18], explicit causal links between two features are not yet identified). In this work, as exact pathologies of AMD are not established, causal graph in Fig 1 was considered as an alternative (domain-based) benchmark causal network when validating or evaluating extracted latent causal variables from retinal fundus images in experiments.

**Fig 1.**
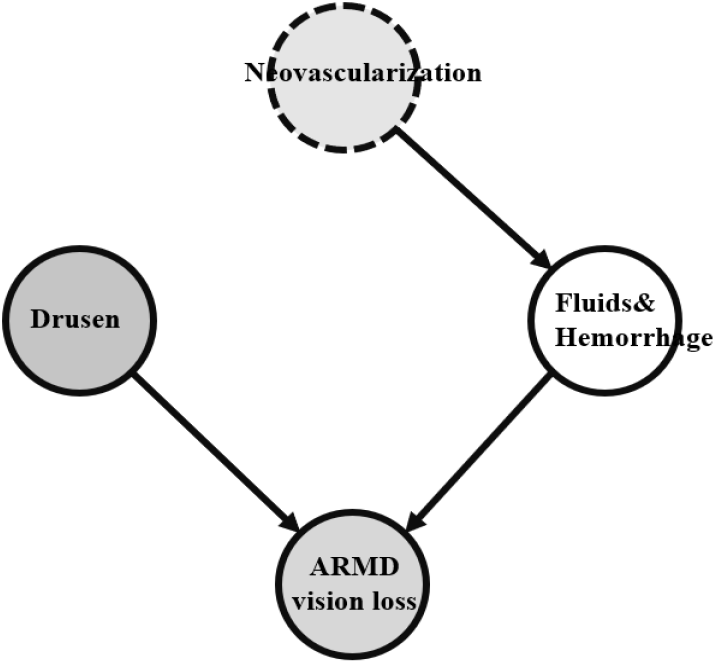
Visualization of Simplified AMD Causal Mechanism.

## B. Related Works

With recent advancements in ML/DL architectures, various applications of AI in AMD analysis have been reported to be successful in diverse environments. Especially for OCT image analysis, CNN based modelling was found successful in most research. For example, using modified VGG16 CNN for 61-line raster macula scan OCT image classification, 92.77% AUROC and 87.63% binary AMD classification accuracy were achieved [12]. In [12], it was also found that pathologic regions can be retrieved from optical coherence tomography image based VGG16, which implies that key features in detecting AMD can be extracted through CNN architecture based deep learning approaches. Furthermore, implementing two CNN based deep residual neural networks: ResNet50 and attention mechanism based ResNet, AMD classification modelling was driven to achieve higher performance in detecting macular degeneration from OCT images, which led to higher classification accuracy when compared to HOG-SVM and VGG [13].

To make classification performance more robust to image preprocessing steps such as RoI extraction and retinal flattening, multi-level dual-attention based CNN approaches were also implemented for advancements in related works [19]. Under Duke and NEH datasets, multi-level DAM achieved higher performance, compared to existing methodologies such as multi-scale CNN ensemble or LACNNs. Apart from focusing on enhancing classification performance, increasing the interpretability of deep learning based AMD classification models was also studied by researchers. For example, under a finely trained VGG16 AMD classification model, gradient-weighted class activation mapping (GradCAM) was implemented as an XAI method to visualize major features that affected the decision of the model [14]. Successfully achieving interpretability, it was found that reliable classification results can be returned under appropriate CNN architectures.

Meanwhile, not only application of Convolutional Neural Networks but also applications of transfer learning in filters or clustering in low dimensional OCT representations were also reported to have significant progress in analyzing AMD [20,21]. For example, Transfer learning based on ImageNet data was implemented for fine-tuning of GoogLeNet in [20], leading to extreme advancements in AMD classification accuracy compared to results from random initialization of filters in normal GoogLeNet based CNNs. Linking expert decisions of ophthalmologists with DNN analysis was also implemented in certain works. In [21], based on a ResNet50 CNN encoder, interpretation of K-means clustering results from low dimensional representations of AMD diagnosed OCT images by retinal specialists was implemented to find novel biomarkers for AMD risk evaluation.

### C. Neural causal DAG extraction

To increase analytic flexibility in conventional causal analysis and to overcome limitations in ML/DL approaches which only guarantee correlation between explanatory variables and target variable, neural causal modeling, which links NNs with Pearl, J’s structural causal model(SCM), was introduced [22-24]. Under the theorem of ℊ -Consistency and *L*_2_-ℊ representation dealt by K, Xia et al. 2022, it was found that DNNs with causal diagram ℊ as a constraint can fully achieve expressivity, while enabling canonical causal inference [22]. That is, when valid causal graph is set as constraint ℊ, non-linear causal effects or relations, which were hard to extract in conventional approaches, can be relatively easily extracted from training neural networks under gradient descent algorithms. Thus, finding valid ℊ became a crucial task to perform successful NCM analysis, which was linked to vast exploitations of algorithms that can extract adjacency matrices or DAGs from observational data. Among recent works, methodologies such as NOTEARS, DAG-GNN and Graph Autoencoder based causal structure extraction were introduced and found to be valid in finding ground truth adjacency matrices from observational data in research [25-27].

Among various methods, considering low-complexity and high performance, this study implemented Graph Autoencoder based DAG extraction methods in [27] to find causal interactions between latent representations from retinal fundus images. Graph AutoEncoders (GAE) are AE based models that extract causal relationships between variables by setting an additional adjacency matrix layer between the encoder and decoder under SCM assumptions. (Though GAE introduced in does not explicitly cover permutation invariance or equivariance of GNNs, the term “graph” is used based on its intrinsic ability to compute message passing and the ability to process causal graphs). In generalized SCM, ground-truth causality for feature set X can be defined as structural equation in (1). Here, any existing DAG relationships can be statistically modeled when *g*_1_, *g*_2_ are assumed as high dimensional functions that can cover both linearity and non-linearity.

As universal approximation theorem holds for deep neural network structures, use of multi-layer perceptrons (MLP) can approximate true *g*_1_, *g*_2_ functions with validity, which implies that under significant convergence in MLPs of Fig 2, true adjacency matrix *A* in causal graph can be successfully returned by setting *A*’s components as trainable weight parameters in neural layers. In GAE, this weight training procedure is computed based on the objective function of (2).

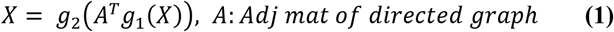

**Fig 2.**
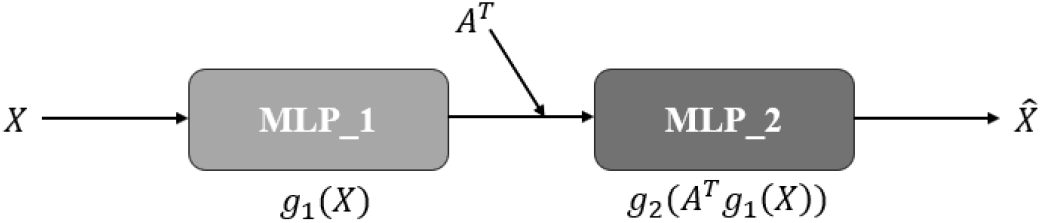
Visualization of Graph AutoEncoder process [27]. Weights are updated under the discrepancy between *X* and 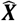.

In the optimization objective of (2), parameters *θ*_1_,*θ*_2_ denote network weights from *MLP*_1_ and *MLP*_2_, *λ*‖*A*‖_1_ denotes *L*_1_ regularization for adjacency matrix 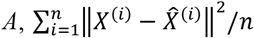 denotes the reconstruction error of a vanilla autoencoder, and *tr*(*e*^*A*⨀*A*^) = *d* works as an equality constraint that constrains weights in *A* to follow the acyclicity assumption of DAG. With Lagrange multiplier set as *α*, optimization problem of (2) can be approximated by augmented Lagrange multiplication of (3). Based optimization of (3), trainable weights in *A, θ*_1_ and *θ*_2_ are updated using gradient descent methods. On the other hand, parameters *α, ρ* are updated under iterative algorithms in (4).

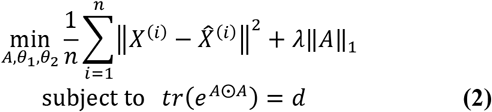

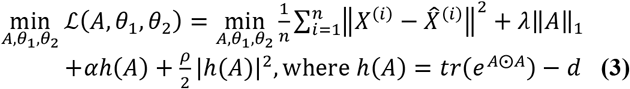

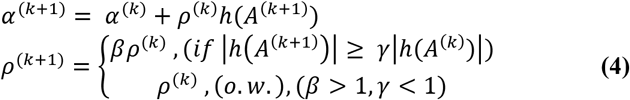

## III. Methodology

### A. Data

For reproducibility, Retinal Fundus Multi-Disease Image public Dataset(RFMiD) was selected for analysis [28]. RFMiD dataset is an open access fundus image dataset acquired from the Eye Clinic of Sushrusha Hospital(India). Consisting of 1920 training images, 640 validation images and 640 test images, each retinal fundus image was taken in examinations from 2009 to 2020 using TOPCON TRC-NW300, TOPCON 3D OCT-2000 and Kowa VX-10. Each set of data was consisted of retinal fundus images and corresponding diagnosis or labels provided by ophthalmology experts. Specifically, each image in RFMiD was first classified into abnormal/normal retina, and then further classified into 45 types of retinal abnormalities or diseases such as presence of diabetic retinopathy, AMD, myopia, etc. As a multiclass retinal disease dataset with diverse quality, RFMiD dataset can highly represent real medical environments where patients can have multiple diseases at the same time and where medical devices can be aged or outdated. Based on its advantages and high reproducibility, many researchers implemented RFMiD in creating CNN models for diverse retinal risk forecasting [29-31].

Focusing on AMD, this research first utilized AMD labels and corresponding images in train data for representation learning (*each image was reshaped to 176×176×3). In RFMiD, AMD was labeled to images if at least one of the following conditions was met: features of neovascular AMD, drusen around the macula, and geographic atrophy related to fovea. Here, to avoid issues from extreme class imbalance (Non AMD: AMD = 1820:100) in training, random under sampling in the majority class was implemented to match the proportion of 200:100. After causal representation learning being completed, AMD patient data and normal retinal fundus data were used to construct high performing AMD classification ML/DL models. With training being complete, AMD and normal retinal fundus image data from RFMiD test dataset was implemented to check prediction performance of fitted ML/DL models.

### B. Latent causal representation learning from fundus image

To find underlying causal interactions from only retinal fundus images with less complexity, explicit latent causal representation learning with graph autoencoder set as the latent DAG search neural algorithm was implemented. Unlike noise encoding approaches in [32,33], explicit modeling encodes causal variable itself for representation. To be specific, latent causal representation learning [32-34] is based on the assumption that when observational space *X* is defined, there exists an unobservable underlying latent mechanism: *Z* →_*g*_ *X*, where latent variables in set *Z* are connected under Pearl, J’s structural causal modeling(SCM) depicted in (5)-(6) [23,24].

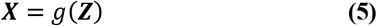

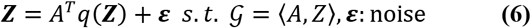

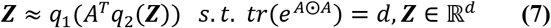

Here, causal graph and the corresponding adjacency matrix are each denoted as 𝒢 and *A*. For causal graph 𝒢 to meet the conditions of DAG, when variables ***Z*** are sorted from parent node to children node, reformulated adjacency matrix *A* should be a triangular matrix with diagonal elements all being 0 to avoid self-loop based edges. For simplicity, these conditions can be replaced by the term in (2), which leads to an alternative definition of (7) to denote a generalized version of latent causal model under DAG 𝒢. In this research, by setting retinal fundus image data as observational variables ***X*** and assuming AMD mechanism components in Fig 1 as ***Z***, causal representation learning was computed as in Fig 3-”Graph Autoencoder part”.

**Fig 3.**
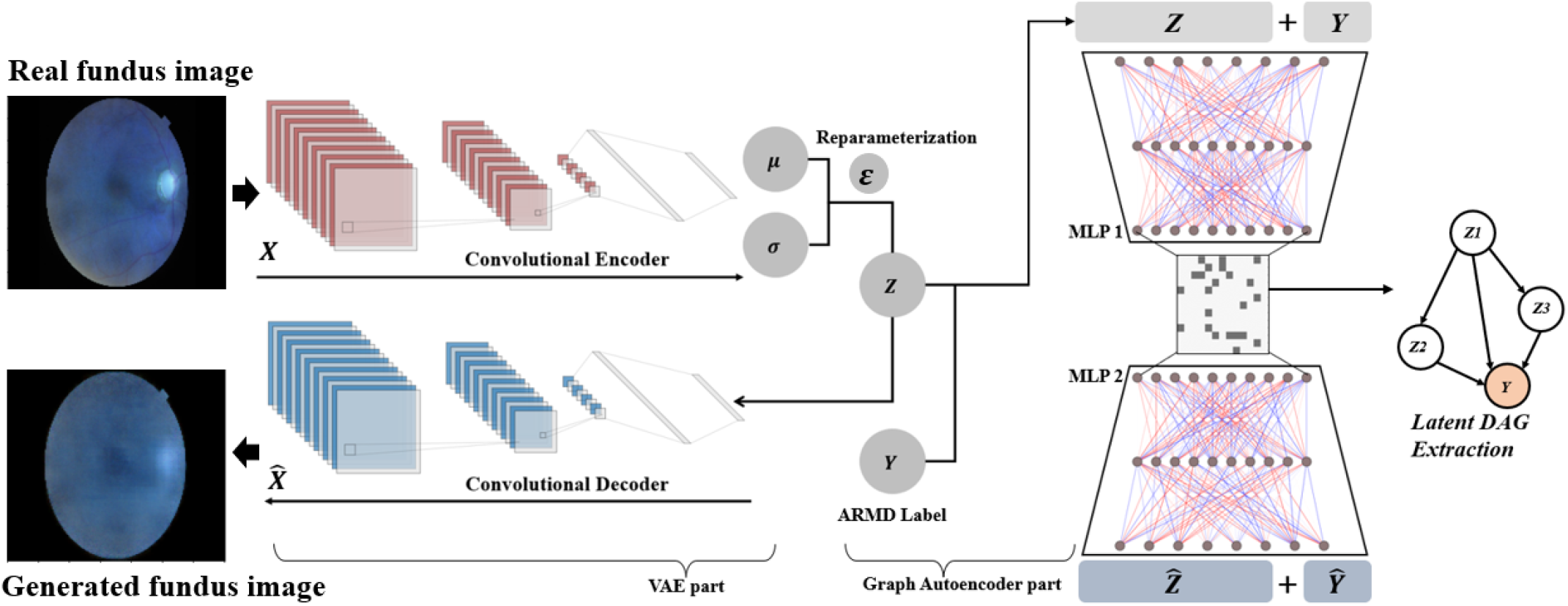
Visualization of CVAE based Causal Representation Learning from RFMiD images.

By first constructing a Convolutional VAE structure to reconstruct retinal fundus images and concatenating AMD label Y with extracted latent representation vector Z, W = ⟨*Z, Y*⟩ was considered as input for Graph Autoencoder based inference. Based on the idea of simultaneous continuous gradient descent for latent causal DAG extraction and VAE fitting in [32], Optimization objective of CVAE weights were defined as in (8). For ℒ_*VAE*_, negative Evidence Lower Bound(ELBO) was used to approximate VAE loss under proof of (9), whereas for ℒ_*GAE*_, optimization problem formation dealt in II.C. was implemented to achieve latent causal DAG *𝒢*. In (9), as KL-divergence is a non-negative value, ELBO can be defined as the lower bound for log-likelihood. Therefore, under an MLE approach, maximization of ELBO loss leads to the maximization of log-likelihood, which makes it possible to set negative ELBO loss values as optimization objectives in VAE [35]. Under *p*(·)∼*Normal*, 1-step (L=1) Monte-Carlo simulation in reparameterization and an identity covariance matrix assumption, ELBO can be further reduced to a mean squared error (MSE) + regularization term form in (10) [35]. Although assuming a Bernoulli distribution for *p*(·) would be ideal for

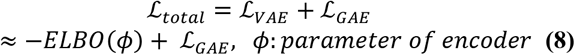

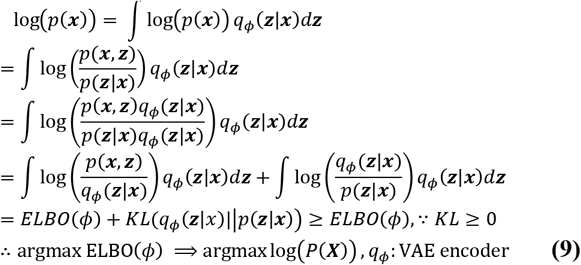

image generation as it leads to cross-entropy loss, due to large discrepancies generated between ℒ_*VAE*_ and ℒ_*GAE*_ under the use of cross-entropy, normal distribution assumptions were chosen to achieve balanced convergence in both objectives. Furthermore, to prevent collapse in image reconstruction tasks, an additional weighting parameter: *ω* was introduced for the regularization term in (10). Therefore, ℒ_*total*_ for CVAE optimization was defined as (11). For updating weights in VAE, loss function of ℒ_*total*_ was implemented. For updating weights

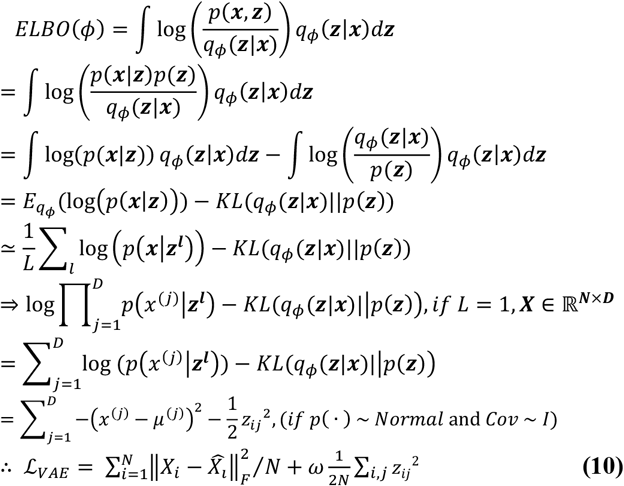

in GAE structures, loss function of ℒ_*GAE*_ was implemented to put more emphasis on valid latent adjacency matrix extraction. Here, in both ℒ_*total*_, ℒ_*GAE*_ definitions, the mean squared error from GAE was given an additional weighting parameter: 𝒱, to ensure balance between sub-objectives: image reconstruction and causal discovery. Thus, latent causal representations ***Z*** that can correspond with underlying features of retinal fundus status were extracted with validity and low complexity.

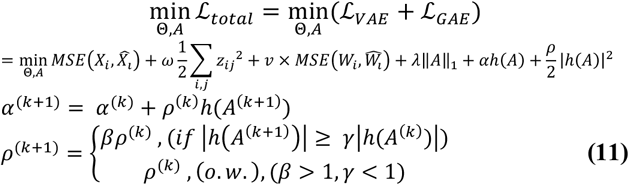

### C. Model Structure Settings-VAE, GAE

Specific structures for explicit causal representation modeling were constructed as follows. For Convolutional VAE’s encoder structures, three convolutional layers and two dense layers after flattening were implemented for latent space retrieval. First convolutional layer was set as having 64 filters with kernel size set as 4×4 and stride size set as 2×2. The second convolutional layer was set as having 64 filters with kernel size set as 6×6 and stride size set as 3×3. The third convolutional layer was set as having 16 filters with kernel size set as 4×4 and stride size set as 2×2. After flattening data from the third layer and adding an additional dense layer with 256 hidden nodes, fully connected dense layer with 2|𝒵| hidden nodes was set for ***µ*** and log (***σ***^2^) in VAE. After generating latent variable ***Z*** using standard gaussian noise ***ε*** based reparameterization: ***µ*** + ***σ*** ⨀ ***ε***, a convolutional decoder which has a symmetrical structure with the former encoder was constructed for retinal image reconstruction. Visualization of VAE structures was computed as in Fig 4. To avoid over-fitting issues, dropout layers with a dropout rate of 10% were implemented between Conv2D or Conv2DTranspose layers. For activation functions, ReLU functions were implemented for conv layers, while ELU functions were implemented for dense layers. Furthermore, for efficient weight initialization, He-Normal initialization [36] was implemented in all hidden layers. All structures for CVAE were constructed based on Python Tensorflow Keras package under tensorflow version 2.19.0.

**Fig 4.**
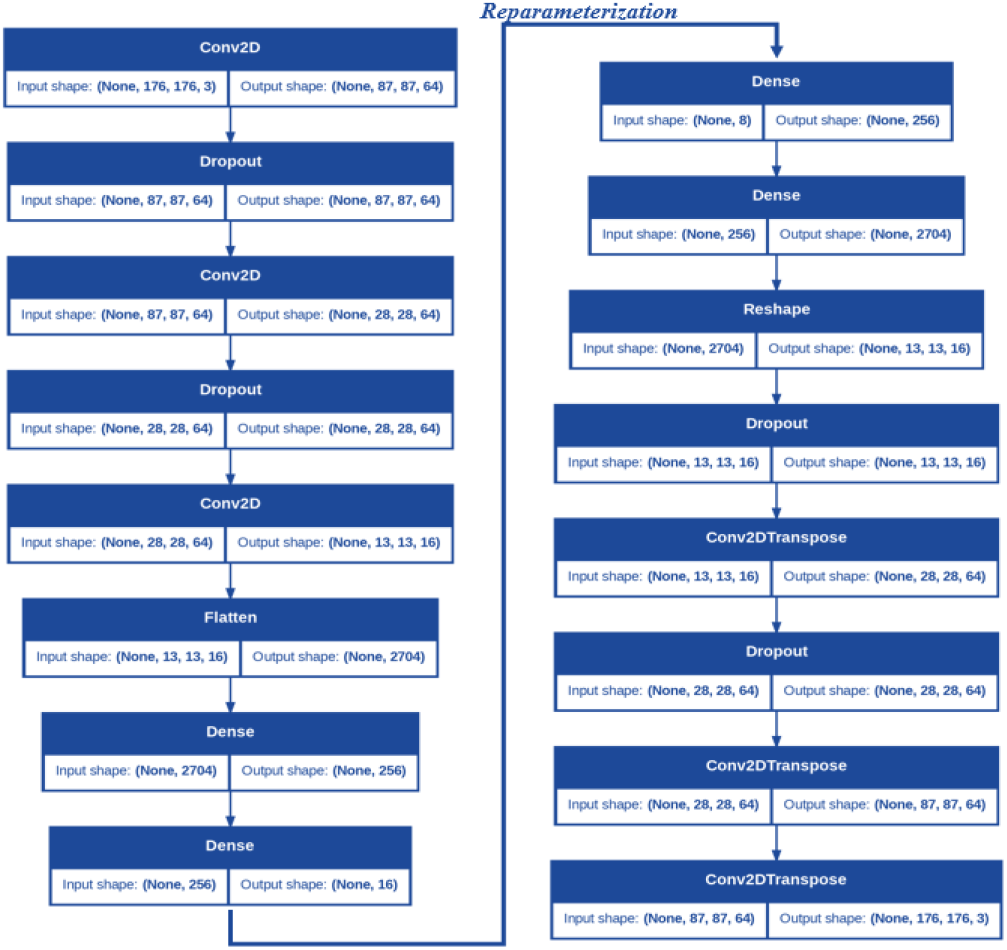
Visualization of CVAE structure framework.

For GAE structures, generalized additive modeling approach within the GAE encoder and decoder was applied to constrain MLPs to follow the form of (7). To perform additive modeling in layers, weight masking based on tensorflow.keras. constraints.Constraint was implemented [37]. Working as a base class for custom constraint generation, tensorflow.keras. constraints.Constraint can be embedded as a kernel constraint in most keras layers with high efficiency. Using weight masking, each input variable in *W* was assigned a separate MLP with two hidden layers, each having 5 hidden nodes, and an output layer, which has only one node. Therefore, a neural network that has 2 non-FCN hidden layers with each having 5(|𝒵| + 1) hidden nodes and an output layer with (|𝒵| + 1) nodes was constructed for both the encoder and the decoder: Fig 5. Apart from basic AE structures, an adjacency weight matrix layer with trainable weights was implemented between two NNs for GAE to extract true edges in latent causal graph *𝒢*. In further steps, weighted adjacency matrix after training was transformed to a binary adjacency matrix using importance based sequential selection.

**Fig 5.**
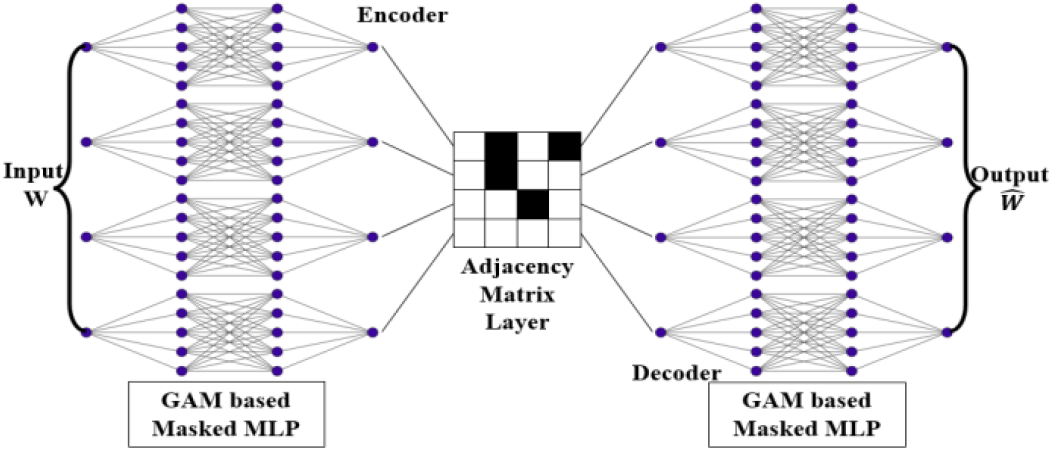
Example of GAE framework with 4 inputs. Each feature was assigned individual MLPs using weight masking.

### D. Experimental Design

Process of training weights in CVAE-GAE and checking performance of latent causal variable extraction were computed as follows. For updating weights, different learning rates were implemented due to non-identical loss function usage. For CVAE, loss function of *ℒ* _*total*_ and an Adam optimizer with learning rate=0.0005 (encoder) and 0.0004 (decoder) were implemented for optimization. For GAE, loss function of *ℒ* _*GAE*_ and an Adam optimizer with learning rate=0.003 were implemented. For parameter settings regarding GAE’s DAG constraint, initial values were set as *α* =0.6, *ρ* =0.1, *γ* =0.9, *β* =1.01 and *λ* =1.0. For hyperparameters: *ω* and 𝒱, each parameter value was set as *ω* = 0.01, 𝒱 = 2.0. In addition, to enforce fast convergence regarding DAG constraint satisfaction, diagonal elements in adjacency matrix and outgoing edges of *Y* were blacklisted by multiplying a binary mask matrix. Using simultaneous updating for all weights, 850 epochs under full-batch gradient descent were computed for optimization.

After training being complete, for convergence check, MSE of CVAE and loss value of CVAE were used as metrics. Under convergence, optimal weighted adjacency matrix was binarized by converting absolute weight values above the 100(1-*q*)^*th*^ percentile as 1 and absolute weights below the threshold as 0. To extract only highly influential weights, hyperparameter *q* was set as *q*=0.16. Based on these results, true causal graph recovery was checked by computing the structural hamming distance(SHD) between causal graph in Fig 1 and the subgraph from extracted DAG. Specifically, to focus only on AMD mechanisms, only edges that are within 2-hop neighbors from *Y* and are directed toward *Y* were considered. Furthermore, under the assumption that extracted causal features are all elements of true causal set, when SHD values can vary under diverse interpretations of topological structures, minimum SHD was selected as the evaluation metric under best possible match. For latent space dimension, |*Z*| = 8 was selected to incorporate both causal and non-causal information within latent space.

To check AMD mechanism identification in latent space, success in achieving causal disentanglement was checked by comparing reconstructed AMD images and original images under modification of individual *z*_*i*_, with other latent variables being fixed. Lastly, after checking one-to-one correspondence between causal variables in set Z and AMD factors in Fig 1. (neovascularization, drusen, hemorrhage and fluids), intervention analysis based on fitted GAE was discussed as an example to illustrate clinical advantages of using causal latent space. (*All experiments were computed under Google Colab’s basic T4 NVIDIA GPU environments).

### E. AMD prediction model construction

Under successful causal disentanglement in latent representations, ML/DL based AMD prediction models were trained to check prediction performance under latent causal variable usage. Using normal patient data and AMD patient data images in RFMiD training dataset, latent representations *Z* were extracted from trained CVAE-encoders, which were then used as input for ML/DL models with label *Y*. Specifically, from ML models, Random Forest, Extra Trees and Gradient Boosting Machines in sklearn.ensemble were implemented, whereas from DL models, a simple DNN with three hidden layers(128,64,8 nodes) was implemented(epochs=500, learning rate=4e-4, batch size=200). Each model was then evaluated using test dataset with five distinct metrics: classification accuracy, weighted F1 score, precision score, recall score and AUC scores. Based on these results, not only causal mechanism comprehension in vision models but also latent causal variable usage for enhanced AMD prediction was checked.

## VI. Experimental results

### A. CVAE and GAE fitting results

Under latent space dimension set as |𝒵| = 8, CVAE and GAE were simultaneously trained for optimal latent causal representation retrieval. Fitting results of CVAE+GAE were deduced as Fig 6. In both neural network structures, stable convergence in ℒ_*VAE*_ and ℒ_*GAE*_ was checked after 850 epochs of training, which implies that both latent causal representation learning and image reconstruction objectives were successfully achieved. Specifically, ℒ_*VAE*_ of CVAE and ℒ_*GAE*_ were computed as 0.0027 and 0.4348.

**Fig 6.**
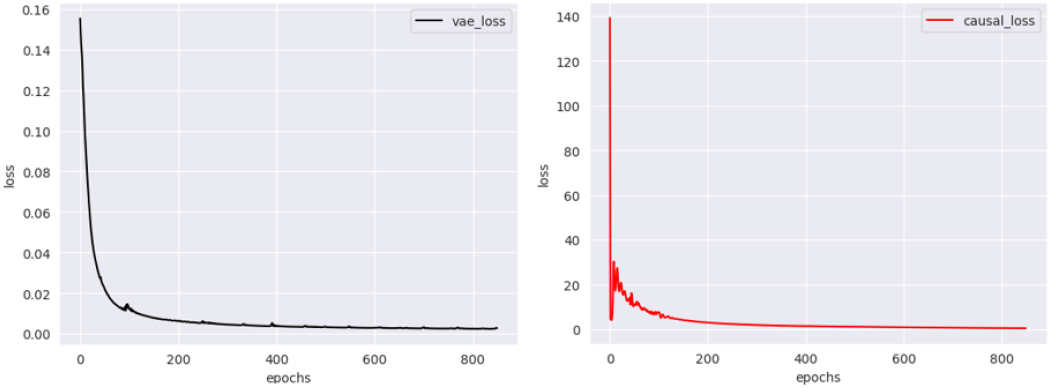
Visualization of latent causal representation learning procedure. (Left) Loss of VAE per epoch (Right) Total loss.

Visual comparisons between original AMD retinal fundus images and generated images were also processed to check validity of training. In Fig 7, 10 images among successful AMD retinal fundus image reconstructions were visualized as examples of CVAE results. Images in the first row denote original retinal fundus images of AMD diagnosed patients, whereas images in the second row denote generated images from CVAE. As it can be seen in Fig 7., abnormally dark light reflections (hypo-reflections), which can be presumed due to retinal fluids or hemorrhage [38,39], and bright clusters, which can be presumed due to AMD related drusen or exudates [38,40], were well captured and reconstructed using latent causal modeling. For example, regarding Case: 5 and Case: 6, CVAE successfully retrieved retinal fundus contaminations (hypo-reflective) and highly reflective legions, which are each presumed to be results of subretinal fluid leakage or hemorrhage and results of drusen or drusen-oriented geographic atrophy around the macular region

**Fig 7.**
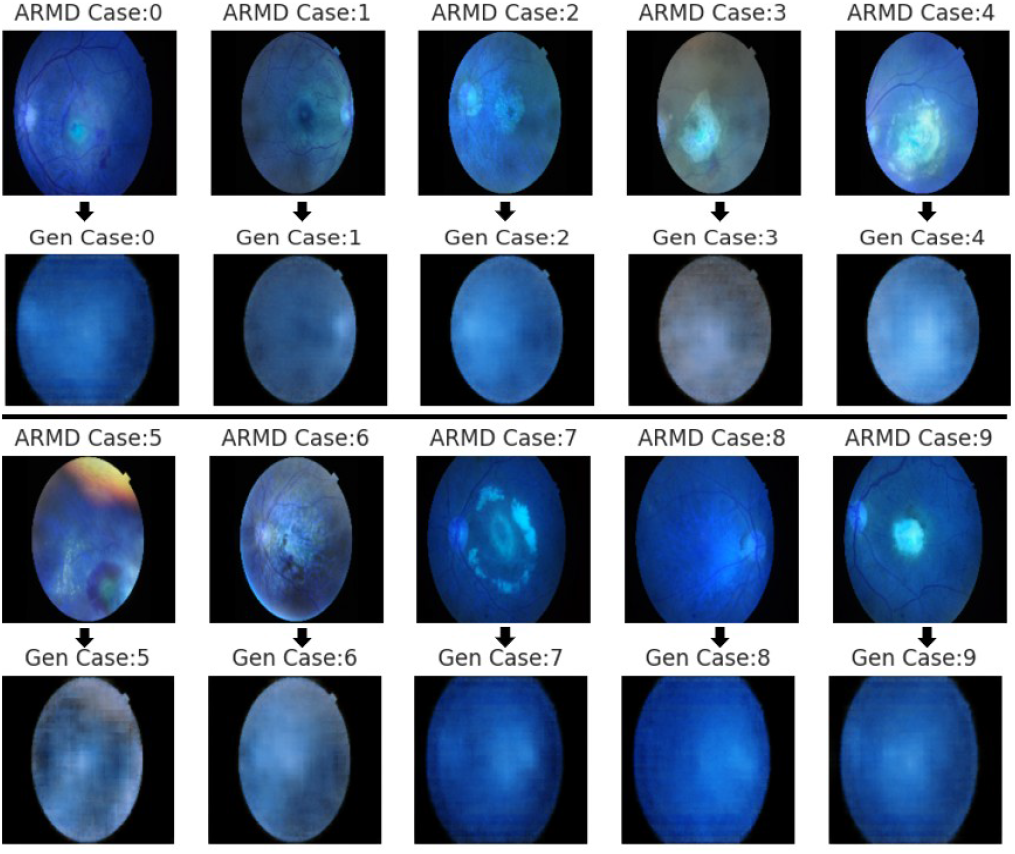
Visualization of CVAE+GAE fitting results. (First row) Original AMD patient retinal fundus images (Second row) Corresponding generated (reconstructed) images.

Based on trained CVAE+GAE structures, weighted adjacency matrix and binarized adjacency matrix from training latent DAGs were also computed as Fig 8. Among latent variables, *z*_0_ and *z*_4_ were found to directly affect AMD status as causal factors. Not having bidirectional edges and having only two sub-cycles within the extracted latent graph, conditions of DAG constraint (directionality, acyclicity) were found to be successfully achieved. In terms of quantitative evaluation, SHD from the extracted subgraph (latent DAG) was computed as 2.0 (Fig 9.), leading to the validation of high possibility of valid AMD mechanism identifications from CVAE+GAE structures. Specifically, among latent causal representations, variables *z*_0_, *z*_4_ and *z*_1_ were found to be possible candidates that can one-to-one correspond with factors in Fig 1. Furthermore, though the extracted causal edge of *z*_4_ to *z*_1_ did not directly match with the assumed domain causal link in Fig 1., regarding recent literatures which stress the importance of drusen as a strong risk factor of CNV [17,18,42], and regarding transition cases of dry AMD to wet AMD, it was found that the extracted causal adjacency matrix has additional potential in terms of considering possible causal links between drusen and neovascularization within the AMD pathology.

**Fig 8.**
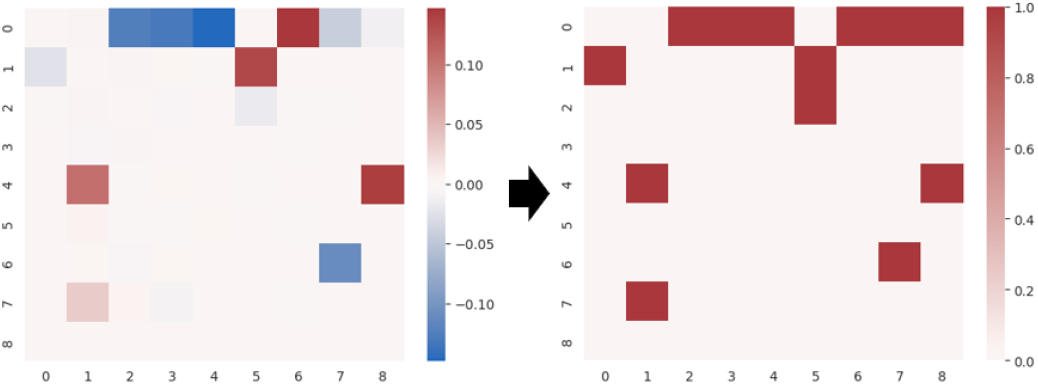
Visualization of trained adjacency matrix from latent causal variables and *Y*: AMD Status. (Right) Binarized *A*.

**Fig 9.**
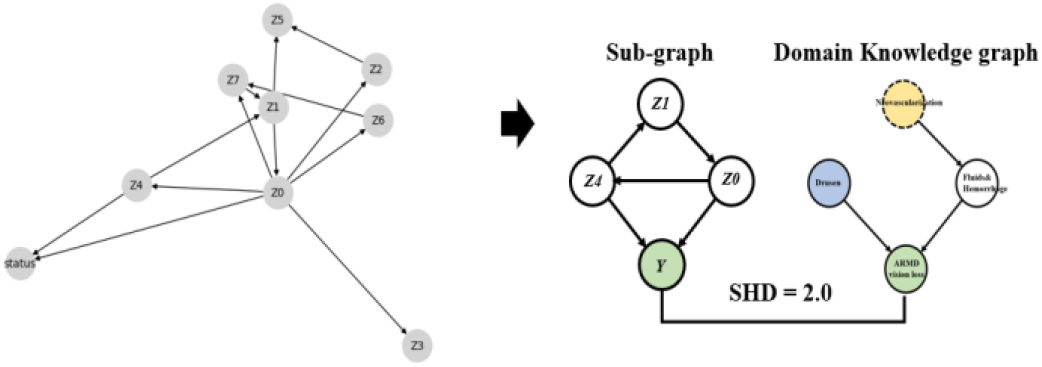
Visualization of extracted latent causal graph and its sub graph. *Y*: AMD status. (Right) Comparison between Domain Knowledge graph and extracted subgraph.

### B. Causal disentanglement check

To validate whether latent causal variables can be matched with domain knowledge-based AMD mechanism factors, causal disentanglement was checked under multiple images by modifying each z within (*z*_0_, *z*_4_, *z*_1_) with other variables being fixed. Here, set (*z*_0_, *z*_4_, *z*_1_) is the encoded latent causal representation of the original retinal fundus image. As retinal status or disease development is not a discrete process, and as no specific segmentations of hemorrhage or drusen were given, qualitative evaluations based on visual characteristics were mainly computed instead of implementing intense quantitative approaches such as entropy-based disentanglement score metrics [41]. Under clinical features or degree of reflectiveness, it was found that *z*_0_ can correspond with hemorrhage and intraretinal fluids, whereas *z*_4_ can correspond with drusen. For example, variations of original retinal fundus images that exhibit representative features of geographic atrophy due to drusen (first original image), hemorrhage (second original image), and both drusen and hemorrhage (third original image) were visualized in Fig 10. In Fig 10, by varying *z*_4_ within -0.4 ∼ 0.8 under equal intervals, only the area that exhibits major signs of drusen was found to be altered. As *z*_4_ values increased, hyper reflectivity was found to be increased (generated) at the corresponding area, which is a representative feature of drusen within retinal fundus images. Furthermore, one-sided Wilcoxon rank sum test for *z*_4_ values from drusen diagnosed and non-diagnosed groups was also computed for quantitative evaluation. Having a test statistic of -2.4309 and p-value of 0.008(< 0.05), latent variable *z*_4_ was also found to be highly related with drusen under statistical significance (significance level=0.05). On the other hand, in Fig 10., by varying *z*_0_ within values: -0.7 ∼ 0.1, only the area which corresponds to low reflectional components or dark matters in the original retina, which are representative features of hemorrhage or intraretinal fluids, was found to be altered. Specifically, as *z*_0_ values deviated from -0.3, both size and darkness of specific regions where existence of hemorrhage is presumed in the original image increased dramatically. (*major changes in the generated retina were emphasized as red circles in Fig 10).

**Fig 10.**
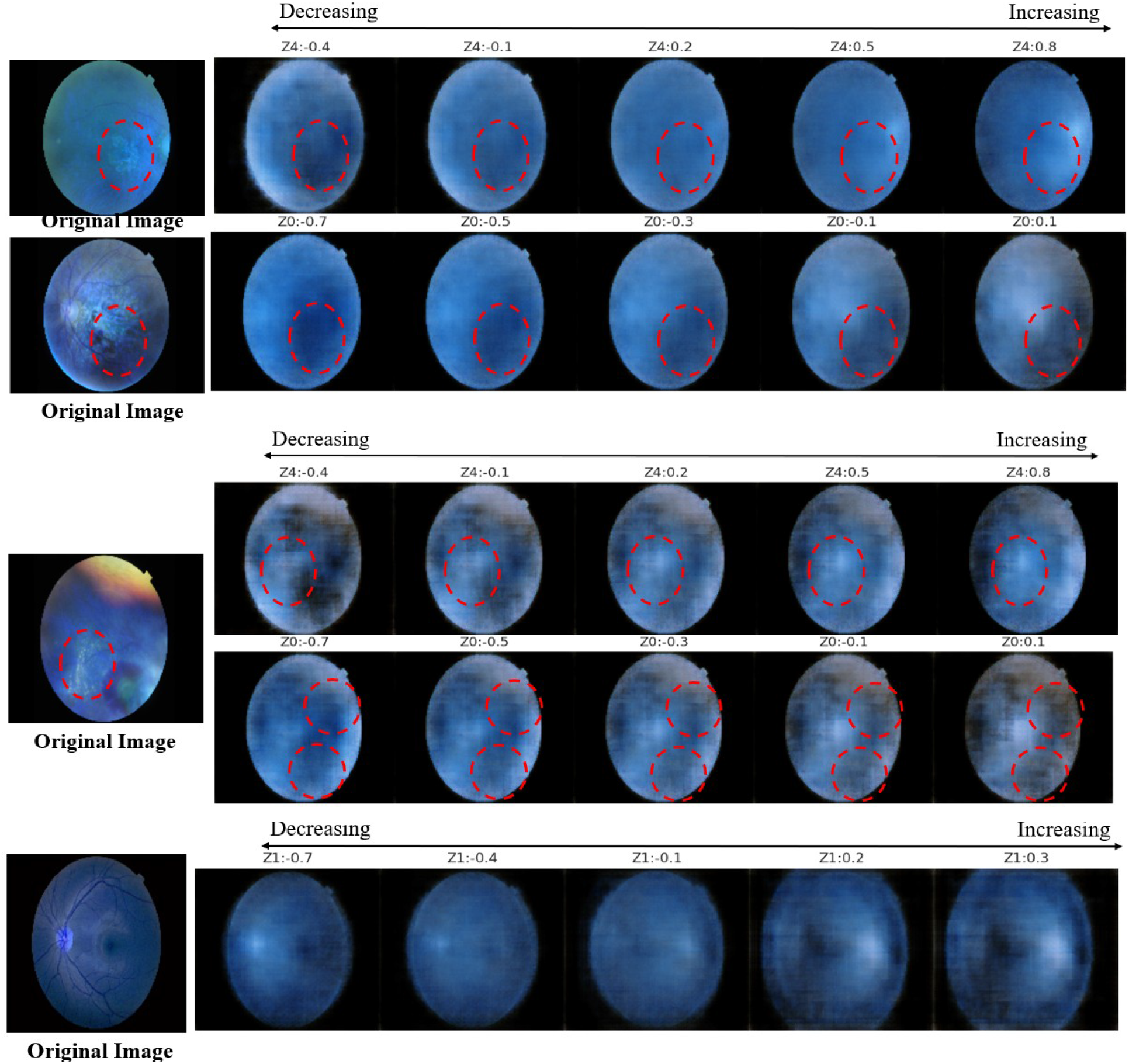
Visualization of Causal disentanglement check on random image. (Right) Original retinal image, (Left) Varying individual latent factors in CVAE reconstruction. Z0 and Z4 were both found to be valid causal representations of factors in AMD mechanism.

On the other hand, *z*_1_ in latent representations exhibited weak correspondence with neovascularization. For example in Fig 10.(fifth row), as *z*_1_ increased from -0.7 to 0.3, a hypo-reflective cluster, which was found to be concentrated around the optic disc region, and the total magnitude of the reconstructed fundus itself were found to be altered. Considering low disentanglement and characteristics of neovascularization, which is hard to identify under fundus images and is found to be prevalent around the macula, it was deduced that *z*_1_ cannot be classified as a proper causal representation of angiogenesis. However, considering the causal linkage between *z*_0_ as a parent node in adjacency matrix of Fig 8., it was found plausible to say that *z*_1_ has possibility of representing certain states of parental variables regarding neovascularization, such as aging or inflammations in retina.

Thus, not only having significant success in extracting valid latent causal space, but also having substantial success in validly identifying major components of AMD, it was deduced that trained CVAE-GAE model can enable accurate retinal fundus analysis under significant comprehension on latent causal structures, and can simulate interventional effects or results of treatments on AMD patients by altering target values among variable set *Z* with reliability. For example, when sufficient regression or ATE analysis between novel treatments and *z*_0_ is driven, clinicians can simulate customized visual results of treatments, such as the progress of intraretinal fluid elimination within the retina, in the AMD risk patient’s retina beforehand by using interventional GAE outputs, which can be acquired from replacing current input *z*_0_ values to estimated 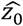 values, in generating individual retinal fundus images.

### C. AMD prediction modeling using latent causal variables

Based on the success in latent causal AMD-related factor extractions, this research attempted to construct a causal feature based AMD diagnosis prediction model using pre-trained CVAE encodings. Explanatory variables for training were computed via encoding Normal and AMD diagnosed patient’s retinal fundus images among original train data into 8-dimensional latent causal features, whereas target variables for training were set as corresponding labels(0: normal, 1: AMD). After training, prediction models were evaluated using test data under identical approach where encodings of images are considered as explanatory variables. Regarding each ML model settings, a Random Forest Classifier(RF) with 100 trees(max depth=10), Gradient Boosting Machine with 100 trees, and an Extra Trees Classifier with 100 trees(max depth=10) were implemented. For DL model settings, a deep neural network with two fully-connected dense layers was implemented. Specifically, for analysis, simple DNN which has 128,64 and 8 hidden nodes for each hidden layer, ELU activation functions for hidden layers, batch normalization between layers and a sigmoid activation function for binary outputs was constructed for experiments. To consider class imbalance in ML model fittings, hyperparameter “class_weight” in sklearn.ensemble models was set as “balanced” in RF and ET. In DL model settings, TensorFlow’s Binary Focal Cross Entropy with class balancing=True, alpha=0.7 and label_smoothing=0.4 was implemented as a loss function to consider class imbalance. Based on above settings, experimental results were deduced as TABLE I (*best metric value in bold). When fitting models to training data, all four models achieved performance in classifying train data. Among models, DNN and RF achieved the highest accuracy score of 100%. In all models, accuracies and F1-scores exceeded 94%. Thus, it was found that using latent causal representations instead of image itself can also lead to successful fitting in AMD classification. Meanwhile, in evaluation based on test data, though achievement of high ROC AUC scores was found in all models, significant variance in other prediction performance metrics was identified among different types of models. Among four models, DNN showed highest prediction performance in most metrics (accuracy: 92.1%, weighted precision: 91.9%, weighted recall: 92.1%, weighted f1-score: 91.9%), whereas GBM showed lowest performance in most metrics(accuracy: 84.85%, weighted precision: 83.34%, weighted recall: 84.85%, weighted f1-score: 82.44%) based on test data.

**TABLE I.** Performance metrics of AMD prediction models (DNN,RF,GBM,ET)

| Model | Train data |  | Test data |  |  |  |  |
| --- | --- | --- | --- | --- | --- | --- | --- |
|  | Accuracy | F1-score | Accuracy | Weighted Precision | Weighted Recall | Weighted F1-score | ROC AUC |
| DNN | **100%** | **100%** | **92.12%** | **91.85%** | **92.12%** | **91.85%** | 87.10% |
| RF | **100%** | **100%** | 86.06% | 86.59% | 86.06% | 83.00% | **91.72%** |
| GBM | 98.00% | 94.74% | 84.85% | 83.34% | 84.85% | 82.44% | 90.66% |
| ExtraTree(ET) | 98.80% | 96.91% | 86.06% | 84.84% | 86.06% | 84.53% | 91.33% |

Unlike ML models, which showed over-fitting result or limitations in extracting true positives (AMD cases) in terms of weighted-recall score and f1-scores, DNN showed consistent performance in all metrics. Specific results of DNN prediction performance were visualized in Fig 11. According to the confusion matrix in Fig 11., though high class imbalance was present in test data, it was found that DNN had substantial success in detecting AMD retinas among mixed data, while minimizing false positive predictions. Having a weighted F1-score of 91.9%, specificity of 97.0% and accuracy of 92.1%, use of latent causal factors extracted from retinal fundus images itself in DL architectures was found to be effective in achieving reliable(robust) and valid AMD predictions. This implies that latent causal modeling based on previous CVAE + GAE structures not only enables valid causal intervention analysis or intervention simulations for specific AMD treatments, but can also enable successful downstream task performance, which can lead to enhancements in state-of-art AMD diagnosis by connecting conventional deep learning based neural network models with latent causal structures.

**Fig 11.**
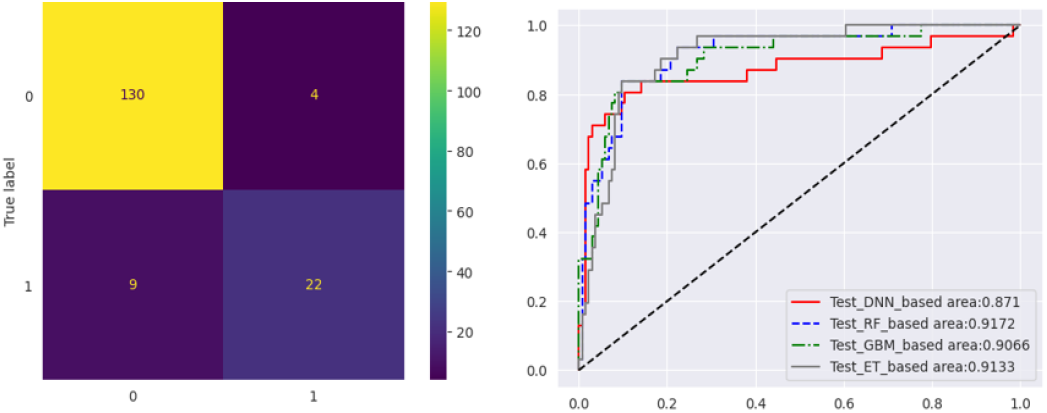
Visualization prediction performance of trained DNN based on latent causal variables extracted from test data. (Right) Confusion Matrix: 0 denotes Normal retinal fundus, 1 denotes AMD retinal fundus. (Left) ROC curves of models.

## V. Conclusion

In this study, for ML/DL models to comprehend or consider the causal mechanism of AMD development in various downstream tasks, Convolutional VAE + Graph autoencoder based explicit latent causal representation learning was implemented to identify causal factors and relations from retinal fundus images directly. Based on successful convergence in both losses: ℒ_*VAE*_, ℒ_*GAE*_, it was deduced that the achieved latent causal graph contains valid underlying causal factors linked to AMD. Furthermore, by first comparing topological structures of extracted graph and pre-defined domain knowledge-based graph under SHD, and then modifying individual latent causal variables within extracted sub-graph, it was found that causal disentanglement or identifications can be fulfilled regarding drusen and NV-based hemorrhage or intraretinal fluids in latent causal representations. Thus, with validity found in CVAE + GAE based latent causal encoders, downstream tasks of classifying and predicting AMD diagnosis were computed based on Normal/AMD diagnosed patient’s retinal fundus images. Results show that though high class imbalance was present, latent causal encoding based DNN achieved extreme prediction performance in most metrics: accuracy = 92.1%, weighted F1-score = 91.9%, specificity = 97.0%. This implies that through the suggested CVAE+GAE model, not only valid causal intervention analysis can be computed with extracted latent causal AMD DAG, but also reliable AMD predictions can be returned while using only low quality retinal fundus images.

Though accurate predictions or treatment (intervention) simulations regarding AMD can be expected using results from this research, there were also clear limitations which should be thoroughly discussed in further works. For example, though hyper/hypo reflective components or contaminations in retinal fundus were well reconstructed in CVAE, reconstruction of specific blood vessels around the macula failed in most image generation cases, which were presumed due to low quality or low resolution in sample images and the intrinsic nature of VAEs. Limitations in implementing quantitative metrics were also found to be present in causal disentanglement checks. Though disentanglement within causal factors were visually apparent under numerous simulations, due to the intrinsic continuity (non-discreteness) in AMD related factors and lack of label data, conventional disentanglement scoring methods (except SHD) were not able to be applied in evaluation. Thus, for further developments, to achieve a more reliable and advanced latent causal analysis regarding mechanisms of AMD, obtaining high resolution fundus image reconstructions using diffusion models or acquiring fully labelled retinal fundus data should be implemented in further research.

